# Radiosensitization of Glioblastoma by the K-ras Inhibitor RMC-6236

**DOI:** 10.64898/2026.05.29.728724

**Authors:** Hong Shik Yun, Tamalee R. Kramp, Mary Sproull, Krishan Thakur, Arnab Chakravarti, Kevin Camphausen

## Abstract

**Purpose:** Glioblastoma (GBM) is characterized by poor clinical outcomes and marked resistance to radiotherapy. Because effective radiosensitizing strategies for GBM remain limited, we investigated whether inhibition of KRAS/RAS signaling could enhance radiation response in GBM. In particular, we evaluated the radiosensitizing potential of RMC-6236, an RAS(ON) multiselective inhibitor that suppresses active RAS signaling across multiple RAS-dependent states.

**Experimental Design:** Human GBM cell lines (U251, LN-18, ACPK1, and OSU61) were treated with radiation, with or without genetic or pharmacological KRAS inhibition. KRAS signaling was suppressed by siRNA-mediated knockdown or RMC-6236 treatment. Radiation-induced KRAS activation and downstream MAPK signaling were assessed by Raf-RBD pull-down assays and immunoblotting. Radiosensitivity was evaluated using clonogenic survival assay. DNA damage persistence, cell cycle distribution, and mitotic catastrophe were analyzed by γH2AX immunofluorescence, flow cytometry, and nuclear morphology assessment, respectively. In vivo therapeutic efficacy was examined in an orthotopic U251 xenograft model.

**Results:** Radiation-induced transient activation and increased KRAS protein expression of KRAS, accompanied by activation of ERK, JNK, and p38 signaling in GBM cells. siKRAS suppressed radiation-induced KRAS and MAPK activation, and significantly enhanced radiosensitivity in all four GBM cell lines. Similarly, RMC-6236 inhibited radiation-induced KRAS activation and attenuated downstream MAPK signaling without reducing the total KRAS protein expression. RMC-6236 significantly increased the radiosensitivity across all GBM cell lines, with dose enhancement factors ranging from 1.33 1.46. Mechanistically, combined treatment with RMC-6236 and radiation increased persistent γH2AX foci and enhanced mitotic catastrophe without producing consistent redistribution of cells into radiosensitive cell cycle phases. In an orthotopic GBM model, the combination of RMC-6236 and radiation significantly prolonged survival compared to that of the control and radiation alone.

**Conclusions:** These findings indicate that radiation-induced KRAS signaling is a functionally important mediator of radioresistance in GBM and demonstrate that inhibition of KRAS/RAS signaling enhances the radiation response *in vitro* and *in vivo*. RMC-6236 may represent a promising radiosensitizing strategy for GBM by suppressing adaptive RAS/MAPK signaling and promoting persistent DNA damage and mitotic catastrophe following irradiation. However, clinical trials of this combination are warranted.

## Introduction

Glioblastoma (GBM) is the most aggressive primary malignant brain tumor in adults and is associated with poor clinical outcomes (1). Because of its infiltrative growth pattern and location within the central nervous system, therapeutic options for GBM are inherently limited, and radiotherapy remains a critical component of treatment (1). However, the efficacy of radiotherapy is substantially limited by intrinsic and acquired radioresistance, which arise through diverse biological mechanisms (2). Together, these features further diminish the therapeutic efficacy of radiotherapy and contribute to the poor prognosis of GBM patients. Accordingly, substantial efforts have been devoted to overcoming radioresistance and improving the therapeutic efficacy of radiotherapy in glioblastoma (2). Among the signaling pathways implicated in radioresistance, the RAS–MAPK axis has emerged as an important regulator of tumor cell survival following radiation. In several solid tumors, particularly lung cancer, oncogenic KRAS has been associated with reduced radiosensitivity through pro-survival and stemness-related signaling, and experimental studies have suggested that KRAS status can influence radiation response (3, 4). Moreover, radiation itself can activate RAS–MAPK signaling, indicating that inducible pathway activation may also contribute to therapeutic resistance, even in the absence of canonical KRAS mutation (3–6). KRAS is frequently altered in human cancers and plays a central role in the regulation of proliferation, survival, and stress adaptation. Its biological importance is particularly well established in pancreatic, colorectal, and non–small cell lung cancers, where KRAS mutations drive malignant progression and therapeutic resistance (7). These observations have led to major advances in the development of KRAS-targeted therapies, most notably mutation-specific inhibitors targeting KRAS G12C. Agents such as Sotorasib and Adagrasib have demonstrated clinically meaningful activity in tumors harboring KRAS G12C mutations by selectively targeting the inactive GDP-bound state of mutant KRAS (8, 9). However, the therapeutic scope of current allele-specific KRAS inhibitors remains largely restricted to tumors harboring the corresponding KRAS mutant alleles. Tumors driven by wild-type KRAS or sustained by upstream receptor tyrosine kinase signaling may evade durable pathway suppression through adaptive reactivation of wild-type RAS signaling (9, 10). These limitations have stimulated an interest in broader RAS-targeted strategies. In preclinical studies, pan-RAS inhibition has shown antitumor activity in multiple RAS-driven cancers, supporting the potential value of targeting both mutant and wild-type RAS signaling (11). Such an approach may be relevant in glioblastoma, where activating KRAS mutations are uncommon (12); however, aberrant RAS–MAPK pathway activation frequently occurs through upstream signaling inputs (13).

RMC-6236 is a RAS(ON) multi-selective inhibitor designed to suppress active RAS signaling across multiple RAS-dependent states rather than targeting a single KRAS mutant allele (13, 14). Preclinical studies and translational modeling have shown that RMC-6236 exerts broad antitumor activity across several RAS-driven cancers, including pancreatic, lung, and colorectal cancers, supporting its potential utility beyond allele-specific KRAS inhibition (13). Given the importance of radiation-induced RAS/MAPK signaling in treatment resistance and the limited availability of effective radiosensitizing strategies targeting this pathway in GBM, we investigated whether KRAS inhibition could enhance radiation response in glioblastoma models. The data presented indicate that RMC-6236 enhances tumor cell radiosensitivity *in vitro* and *in vivo*, which correlates with changes in KRAS downstream signaling. Moreover, sensitization was correlated with delayed dispersion of γ-H2AX and an increase in mitotic catastrophe. Most importantly, the drug was effective in an intracranial tumor model, demonstrating its ability to cross the BBB.

## Materials and Methods

### Cell lines and treatments

The human glioblastoma cell lines U251, LN-18, ACPK1, and OSU61 were used in this study. All cell lines were maintained in DMEM supplemented with 10% fetal bovine serum (Invitrogen) at 37 °C in a humidified atmosphere containing 5% CO₂. The primary glioblastoma cell lines ACPK1 and OSU61, derived from patient tumor specimens, were authenticated by a neuropathologist at Ohio State University. For in vitro experiments, cells were treated with the KRAS inhibitor RMC-6236 (Selleckchem) at concentrations of 5–20 nM for 4 h prior to irradiation unless otherwise specified. RMC-6236 was dissolved in dimethyl sulfoxide (DMSO; Sigma-Aldrich) and diluted in the culture medium immediately before use. Control cells were treated with an equivalent volume of dimethyl sulfoxide (DMSO). The cells were irradiated as monolayer cultures using an XRad-320 X-ray irradiator (Precision X-Ray Inc.) at a dose rate of 2.5 Gy/min. For the radiosensitization experiments, cells were pretreated with RMC-6236 for 4 h before exposure to the indicated doses of radiation (1–8 Gy). Control cells were sham-irradiated under identical conditions without radiation exposure.

### Ras activation assay

Ras activation was evaluated using a Ras Activation Assay Biochem Kit (Cat. #BK008; Cytoskeleton Inc.) according to the manufacturer’s instructions. The assay is based on selective binding of the Raf-1 Ras-binding domain (RBD) to the GTP-bound active form of Ras, allowing affinity purification of activated Ras from cell lysates. Briefly, the cells were treated with RMC-6236 with or without irradiation under the indicated experimental conditions and rapidly washed with ice-cold PBS. Cells were lysed in ice-cold cell lysis buffer supplemented with protease inhibitors provided in the kit, and the lysates were clarified by centrifugation at 4 °C. Protein concentrations were determined and equalized prior to the pull-down assay. Equivalent amounts of total protein were incubated with Raf-RBD agarose beads at 4 °C with gentle rotation to selectively capture the GTP-bound Ras. The beads were then washed with wash buffer and resuspended in the Laemmli sample buffer. Bound proteins were separated by SDS-PAGE and subjected to immunoblot analysis using a pan-Ras antibody supplied in the kit. The level of active Ras (Ras-GTP) was determined by immunoblotting and compared across treatment groups. Total Ras in whole-cell lysates was analyzed in parallel to normalize Ras activation levels.

### Clonogenic survival assay

For clonogenic survival assays, U251, LN-18, ACPK1, and OSU61 cells were seeded into six-well plates at a density of 50–3,200 cells per well, depending on the radiation dose. After attachment for 16 h, the cells were pretreated with RMC-6236 for 4 h and exposed to 1–8 Gy radiation. At 24 h after irradiation, the drug-containing medium was removed and replaced with fresh drug-free medium. The colonies were allowed to grow for 10–14 days, fixed, and stained with 0.1% crystal violet. Colonies consisting of 50 or more cells were scored as survivors and the surviving fractions were calculated. The data represent the mean ± SE of at least three independent experiments.

### Reverse transcription PCR and quantitative real-time PCR

Total RNA was isolated using the Direct-zol RNA Miniprep Kit (Zymo Research), according to the manufacturer’s instructions. Reverse transcription was performed using ImProm-II Reverse Transcription System (Promega) under the following conditions: 5 min at 25 °C for primer annealing, 60 min at 42 °C for cDNA synthesis, and 15 min at 70 °C for enzyme inactivation. Quantitative real-time PCR (qRT-PCR) was performed using the TaqMan Universal Master Mix II with UNG (Thermo Fisher Scientific) and human KRAS TaqMan probes (Thermo Fisher Scientific). Amplification reactions were performed according to the manufacturer’s protocol, and KRAS mRNA levels were normalized to GAPDH as an internal control.

### Transient transfection of small interfering RNA

Small interfering RNAs (siRNAs) targeting human KRAS were purchased from Qiagen. Two independent siRNAs, targeting different KRAS sequences, were used in this study. Cells were transfected with 25 nM siRNA in serum-free medium using Lipofectamine™ RNAiMAX (Thermo Fisher Scientific), according to the manufacturer’s instructions. Following overnight incubation, the medium was replaced with a complete growth medium, and the cells were further cultured for 48 h at 37 °C. The efficiency of KRAS knockdown was evaluated using immunoblot analysis, and the siRNA with the highest knockdown efficiency was selected for subsequent experiments.

### Flow cytometric analysis of cell cycle

Flow cytometry was used to evaluate the cell cycle phase distribution. The cells were treated with RMC-6236 (10 nM) alone or in combination with radiation under the same experimental conditions as those described for the clonogenic assays. For combination treatment, cells were pretreated with RMC-6236 for 4 h prior to irradiation (4 Gy). Following irradiation, the cells were harvested at the indicated time points, fixed with 70% ethanol, and stained with propidium iodide (Sigma-Aldrich). DNA content for cell cycle analysis was determined using an LSR Fortessa™ flow cytometer (BD Biosciences).

### Immunoblot analysis

Whole-cell pellets of U251, LN-18, ACPK1, and OSU61 cells were collected for protein extraction in radioimmunoprecipitation assay (RIPA) lysis buffer (Thermo Fisher Scientific). Total protein concentrations were determined using a BCA protein assay (Thermo Fisher Scientific). Proteins were separated using SDS-PAGE (Mini-Protean TGX™ gels; Bio-Rad) and transferred onto nitrocellulose membranes (Bio-Rad). Membranes were probed with antibodies against KRAS (Invitrogen), ERK1/2, phospho-ERK1/2, p38, phospho-p38, JNK, and phospho-JNK ( Cell Signaling Technology). β-Actin (Cell Signaling Technology) was used as a loading control. Protein bands were visualized using IR Dye secondary antibodies (LI-COR) and quantified with an Odyssey CLx imaging system (LI-COR).

### Immunofluorescent staining for γH2AX foci and mitotic catastrophe detection

Visualization of γH2AX foci and mitotic catastrophe was performed as described previously (15). U251, LN-18, ACPK1, and OSU61 cells were grown on 18 × 18-mm cover glasses and treated with RMC-6236 (10 nM) alone or in combination with radiation (4 Gy). For combination treatment, cells were pretreated with RMC-6236 for 4 h prior to irradiation. Cells were fixed and stained for γH2AX, and images were acquired using a Zeiss upright fluorescence microscope. γH2AX foci were quantified by counting at least 50 cells per experimental condition, and the images were analyzed using the ImageJ software. Nuclear morphology was examined to analyze mitotic catastrophe and nuclear fragmentation was defined as the presence of two or more distinct lobes within a single cell. Mitotic catastrophe was scored for at least 100 cells per condition.

### Orthotopic *in vivo* experiments

For *in vivo* studies, U251 glioblastoma cells engineered to express luciferase and GFP (Lentivirus, LVpFUGW-UbC-ffLuc2-eGFP2) were orthotopically implanted into the right striatum of 8–10-week-old athymic female nude mice (nu/nu; NCI Animal Production Program). Tumor growth was monitored using bioluminescence imaging (BLI) and local irradiation was performed as previously described (16). Six days after tumor implantation, BLI signals were detected in all animals, and the mice were randomized into four treatment groups: control (DMSO), RMC-6236 alone, radiation alone, and combination treatment (RMC-6236 + radiation). Each group consisted of five mice. Mice received 5% DMSO and 95% corn oil or RMC-6236 once daily for three consecutive days, and tumors were locally irradiated following drug administration. The survival of tumor-bearing mice was monitored daily until the onset of neurological symptoms, and the animals were euthanized at the onset of morbidity. Kaplan–Meier survival curves were generated to evaluate the treatment effects. All animal procedures were performed in accordance with the of NIH Guide for the Care and Use of Laboratory Animals and approved by the appropriate Institutional Animal Care Committee. Survival analyses were performed using the GraphPad Prism 8.

## Results

### Radiation induces activation and increased protein expression of KRAS in GBM cells

Previous studies have shown that radiation can activate RAS signaling and downstream MAPK pathways in both cancer and normal cells (6). To determine whether radiation regulates KRAS signaling in GBM, four human GBM cell lines (U251, LN-18, ACPK1, and OSU61) were exposed to 4 Gy irradiation and analyzed at the indicated time points. As shown in Fig. 1A–D (left panels), irradiation induced a transient increase in KRAS protein expression in all four GBM cell lines. This effect was observed in U251 (Fig. 1A), LN-18 (Fig. 1B), ACPK1 (Fig. 1C), and OSU61 (Fig. 1D) cells, with KRAS protein levels increasing at early time points following irradiation and gradually declining thereafter. Concomitantly, irradiation also induced the phosphorylation of downstream MAPK signaling components, including ERK, p38, and JNK. Consistent with these findings, Raf-RBD pull-down assays demonstrated increased KRAS activity following irradiation in all four GBM cell lines (Fig. 1A–D, right panels). These results indicate that radiation activates KRAS signaling and the downstream MAPK pathways in GBM cells. To further investigate the mechanism underlying the radiation-induced changes in KRAS protein expression, we examined KRAS mRNA levels following irradiation. Quantitative RT-PCR analysis showed that KRAS mRNA expression remained largely unchanged after radiation exposure in all four GBM cell lines (Supplementary Fig. S1-1), suggesting that the observed increase in KRAS protein levels was not transcriptionally regulated. CHX chase experiments were performed to determine whether radiation affects KRAS protein stability, cycloheximide (CHX) chase experiments were performed. All cell lines were pretreated with CHX (200 or 500 μmol/L) for 4 h to inhibit de novo protein synthesis followed by irradiation. Under these conditions, there was no increase in KRAS 2 or 8h after radiation exposure (Supplementary Fig. S1-2), suggesting that radiation may regulate KRAS protein expression via a post-transcriptional mechanism. Together, these results demonstrate that radiation induces transient activation and increased protein expression of KRAS in GBM cells, accompanied by the activation of downstream MAPK signaling pathways.

**Figure 1.**
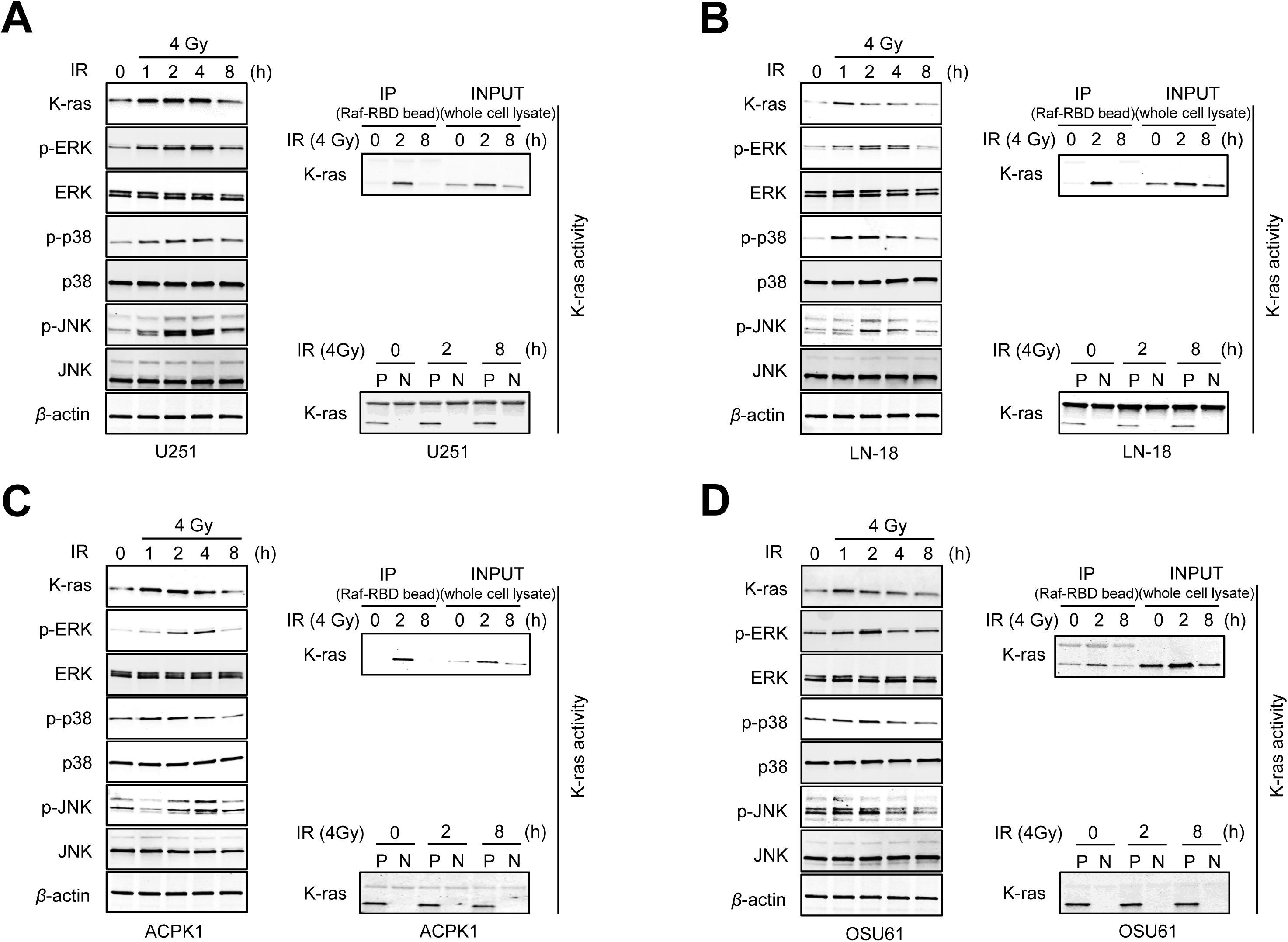
Radiation induces KRAS activation and MAPK signaling in human GBM cells. (A–D) Human glioblastoma (GBM) cell lines U251 (A), LN-18 (B), ACPK1 (C), and OSU61 (D) were exposed to 4 Gy radiation, and cells were harvested at the indicated time points following irradiation. Left panels: Protein levels of KRAS, phosphorylated and total ERK, p38, and JNK were analyzed by immunoblotting in each cell line. β-actin was used as the loading control. Right panels: KRAS activity was assessed using a Raf-RBD pull-down assay followed by immunoblotting for KRAS. Input samples are shown as controls for the total KRAS expression. For the activity assay, P and N denote the positive and negative controls, respectively.

### siKRAS suppresses radiation-induced activation of KRAS and downstream MAPK signaling in GBM cells

To determine whether KRAS mediates radiation-induced activation of MAPK signaling in glioblastoma (GBM), KRAS expression was suppressed by siRNA-mediated knockdown in four human GBM cell lines (U251, LN-18, ACPK1, and OSU61). Cells were transfected with si KRAS (25 nmol/L) and subsequently exposed to 4 Gy radiation, followed by the analysis of KRAS signaling components. As shown in Fig. 2A–D (left panels), siKRAS transfection markedly reduced the basal KRAS protein expression in all four GBM cell lines. In control cells, irradiation induced a transient increase in KRAS protein expression and phosphorylation of downstream MAPK signaling components, including ERK, JNK, and p38. However, siKRAS knockdown markedly attenuated radiation-induced MAPK activation in U251 (Fig. 2A), LN-18 (Fig. 2B), ACPK1 (Fig. 2C), and OSU61 cells (Fig. 2D). Consistent with these findings, Raf-RBD pull-down assays demonstrated that si-KRAS effectively suppressed radiation-induced KRAS activity in all four GBM cell lines (Fig. 2A–D, right panels). Together, these results indicate that KRAS plays a critical role in mediating radiation-induced activation of MAPK signaling in GBM cells and that suppression of KRAS effectively blocks this radiation-induced signaling cascade.

**Figure 2.**
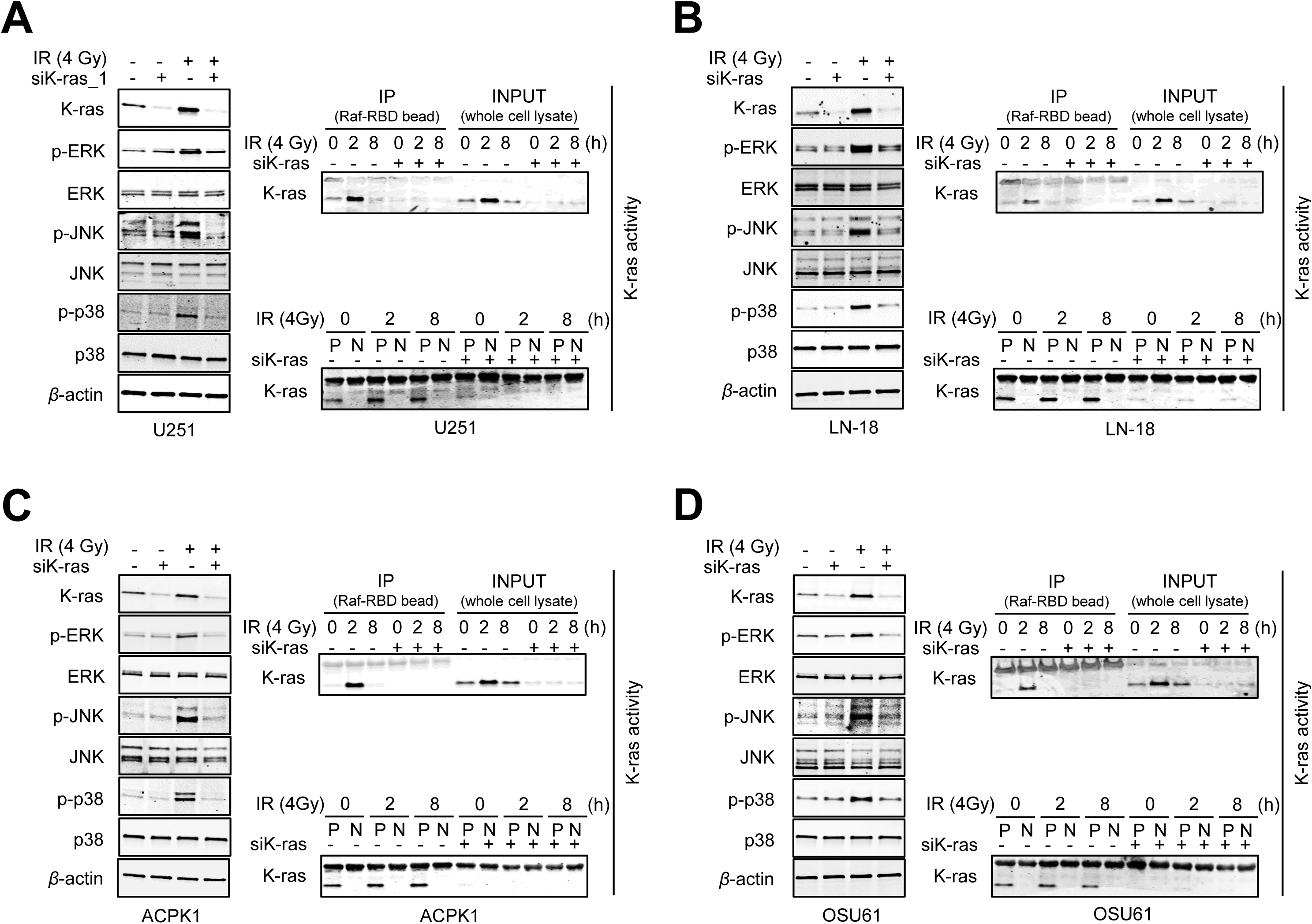
siKRAS suppresses radiation-induced activation of KRAS and MAPK signaling in GBM cells. (A–D) Human glioblastoma (GBM) cell lines U251 (A), LN-18 (B), ACPK1 (C), and OSU61 (D) were transfected with siKRAS (25 nmol/L) and subsequently exposed to 4 Gy radiation. The cells were harvested at the indicated time points following irradiation. Left panels: Protein levels of KRAS, phosphorylated and total ERK, JNK, and p38 were analyzed by immunoblotting in each cell line. β-actin was used as the loading control. Right panels: KRAS activity was evaluated using a Raf-RBD pull-down assay followed by immunoblotting for KRAS. The input samples represent total cellular KRAS levels. For the KRAS activity assay, P and N indicate positive and negative controls, respectively.

### Suppression of KRAS enhances radiosensitivity in GBM cells

Previous studies have reported that inhibition of KRAS signaling can enhance radiosensitivity in cancer cells (4, 5). Consistent with these observations and our findings that KRAS mediates radiation-induced MAPK activation (Fig. 2), we next examined whether KRAS suppression affects the radiation response of glioblastoma (GBM) cells. To address this, clonogenic survival assays were performed following siKRAS-mediated knockdown. Four human GBM cell lines (U251, LN-18, ACPK1, and OSU61) were transfected with siKRAS (25 nM) and subsequently exposed to increasing doses of radiation. As shown in Fig. 3, KRAS knockdown significantly enhanced the radiosensitivity of all the GBM cell lines examined. Compared with control siRNA–transfected cells, siKRAS-treated cells exhibited markedly reduced clonogenic survival following irradiation, with dose enhancement factors (DEFs) calculated at a surviving fraction of 0.1 (SF0.1) of 1.44 U251, 1.31 for LN-18, 1.37 ACPK1, and 1.33 for OSU61, indicating that suppression of KRAS increases radiation sensitivity in GBM cells. Taken together with the results in Fig. 2, these findings demonstrate that KRAS activity contributes to radiation resistance in GBM cells and that inhibition of KRAS enhances cellular sensitivity to radiation.

**Figure 3.**
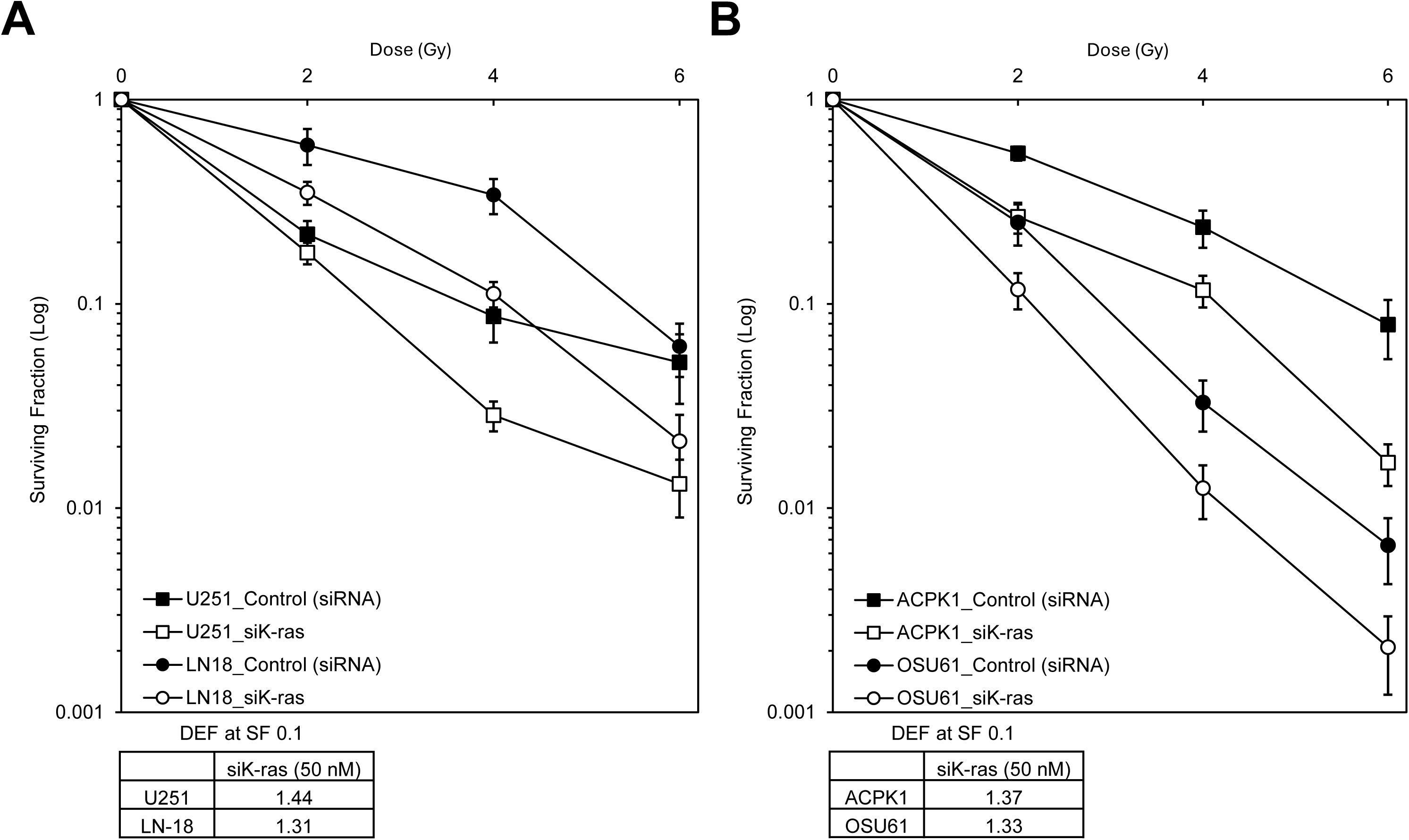
KRAS suppression enhances radiosensitivity in GBM cells. (A-B) Clonogenic survival assays were performed to evaluate the radiosensitizing effect of siKRAS in human glioblastoma (GBM) cell lines. Cells were transfected with siKRAS (25 nmol/L) and subsequently exposed to increasing doses of radiation. (A) Survival curves for U251 and LN-18 cells following transfection with siKRAS or control siRNA and irradiation at the indicated doses. (B) Survival curves for ACPK1 and OSU61 cells following transfection with siKRAS or control siRNA and irradiation at the indicated doses. The dose enhancement factor (DEF) was calculated at a surviving fraction of 0.1 (SF0.1). The DEF values were 1.44 for U251, 1.31 for LN-18, 1.37 for ACPK1, and 1.33 for OSU61. Data represent mean ± SD from independent experiments. Points, mean; bar, SE. DEF, dose enhancement factor; IR, irradiation; SF, surviving fraction.

### RMC-6236 suppresses radiation-induced KRAS activation and downstream MAPK signaling in GBM cells

To determine whether pharmacological inhibition of KRAS suppresses radiation-induced KRAS signaling, four human glioblastoma (GBM) cell lines (U251, LN-18, ACPK1, and OSU61) were treated with RMC-6236, a pan-RAS inhibitor that suppresses both mutant and wild-type KRAS signaling and inhibits downstream MAPK pathway activation (13). Cells were pretreated with RMC-6236 (5, 10, or 20 nmol/L) for 4 h prior to exposure to 4 Gy radiation, and samples were collected 0, 2, and 8 h after irradiation. As shown in Fig. 4A–D (left panels), radiation exposure induced the activation of downstream MAPK signaling pathways, including phosphorylation of ERK, JNK, and p38, in all four GBM cell lines (U251, LN-18, ACPK1, and OSU61). Pretreatment with RMC-6236 attenuates radiation-induced MAPK activation in a dose-dependent manner. Consistent with the mechanism of action of RMC-6236, KRAS protein levels were not significantly altered by drug treatment, in contrast to the reduction observed following si KRAS knockdown (Fig. 2), indicating that RMC-6236 did not reduce KRAS protein expression. To determine whether RMC-6236 inhibited KRAS activity, Raf-RBD pull-down assays were performed. As shown in Fig. 4A–D (right panels), radiation-induced KRAS activity was markedly suppressed by RMC-6236 treatment in all four GBM cell lines. Together, these findings demonstrate that pharmacological inhibition of KRAS with RMC-6236 suppresses radiation-induced KRAS activation and downstream MAPK signaling in GBM cells.

**Figure 4.**
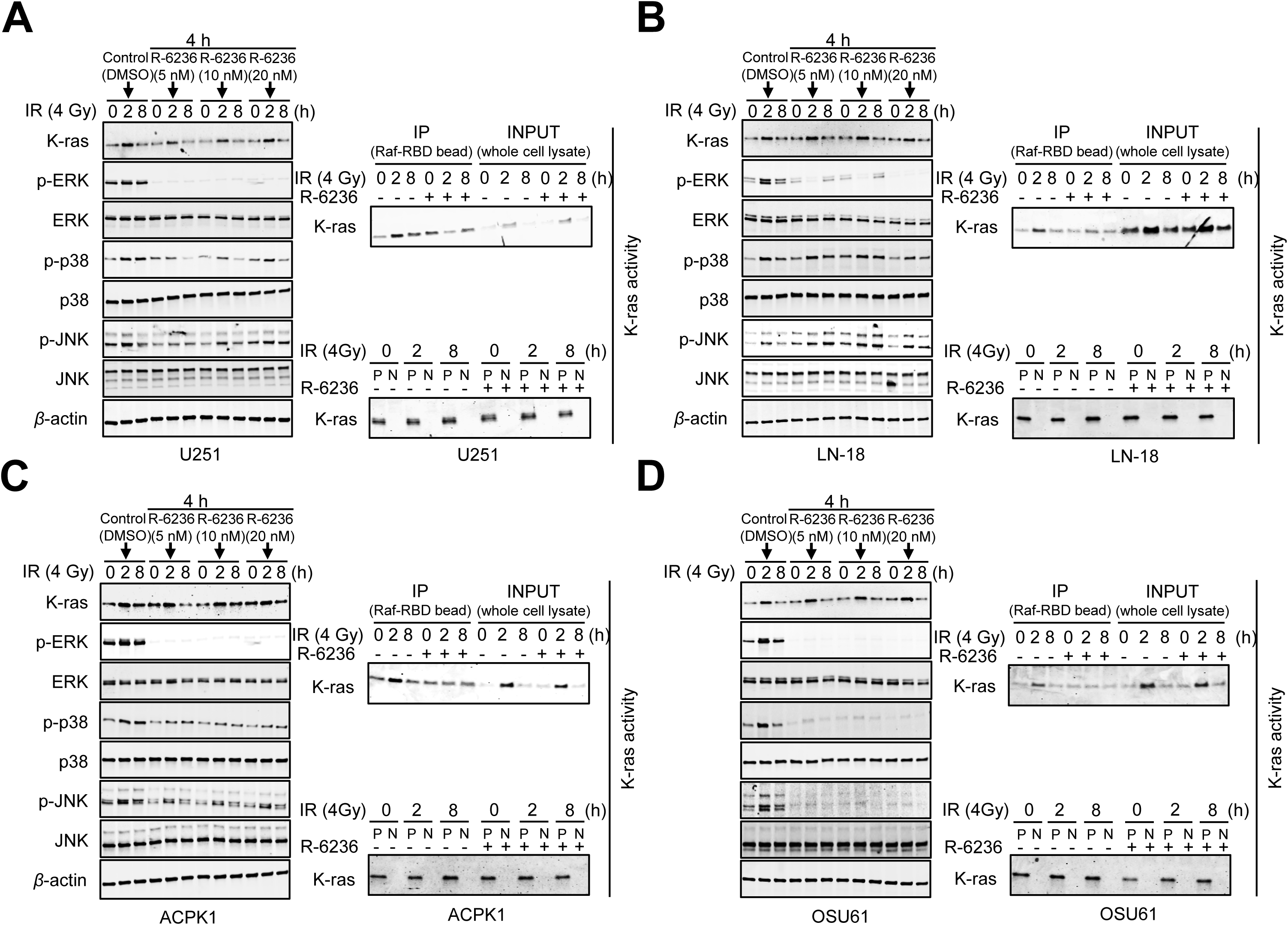
RMC-6236 suppresses radiation-induced KRAS activation and MAPK signaling in GBM cells. (A–D) Human glioblastoma (GBM) cell lines U251 (A), LN-18 (B), ACPK1 (C), and OSU61 (D) were pretreated with RMC-6236 (5, 10, or 20 nmol/L) for 4 h and subsequently exposed to 4 Gy radiation. The cells were harvested at the indicated time points (0, 2, and 8 h) following irradiation. Left panels: Immunoblot analysis showing the protein levels of KRAS, phosphorylated and total ERK, JNK, and p38. β-actin was used as the loading control. Right panels: KRAS activity assays were performed using Raf-RBD pull-down assays followed by immunoblotting for KRAS. The input samples represent total cellular KRAS levels. P and N indicate the positive and negative controls, respectively.

### RMC-6236 enhances radiosensitivity in GBM cells

Given that RMC-6236 effectively suppressed radiation-induced KRAS activation and downstream MAPK signaling in GBM cells (Fig. 4), we examined whether pharmacological inhibition of KRAS enhanced the radiation response of GBM cells. To address this, clonogenic survival assays were performed using four human glioblastoma (GBM) cell lines (U251, LN-18, ACPK1, and OSU61). The cells were pretreated with the KRAS inhibitor RMC-6236 (10 nmol/L) for 4 h prior to exposure to increasing doses of radiation. As shown in Fig. 5A, treatment with RMC-6236 significantly enhanced radiosensitivity in U251 and LN-18 cells, as evidenced by a reduction in clonogenic survival compared to DMSO-treated control cells. Similar results were observed in ACPK1 and OSU61 cells (Fig. 5B), in which RMC-6236 treatment markedly decreased cell survival following irradiation. The DEF values were 1.39 for U251, 1.34 for LN-18, 1.46 for ACPK1, and 1.33 for OSU61, indicating that pharmacologic inhibition of KRAS increases radiation sensitivity in GBM cells. Together, these findings demonstrate that RMC-6236 enhances the radiosensitivity of GBM cells, which is consistent with the effects observed following si-KRAS-mediated suppression of KRAS signaling (Fig. 3).

**Figure 5.**
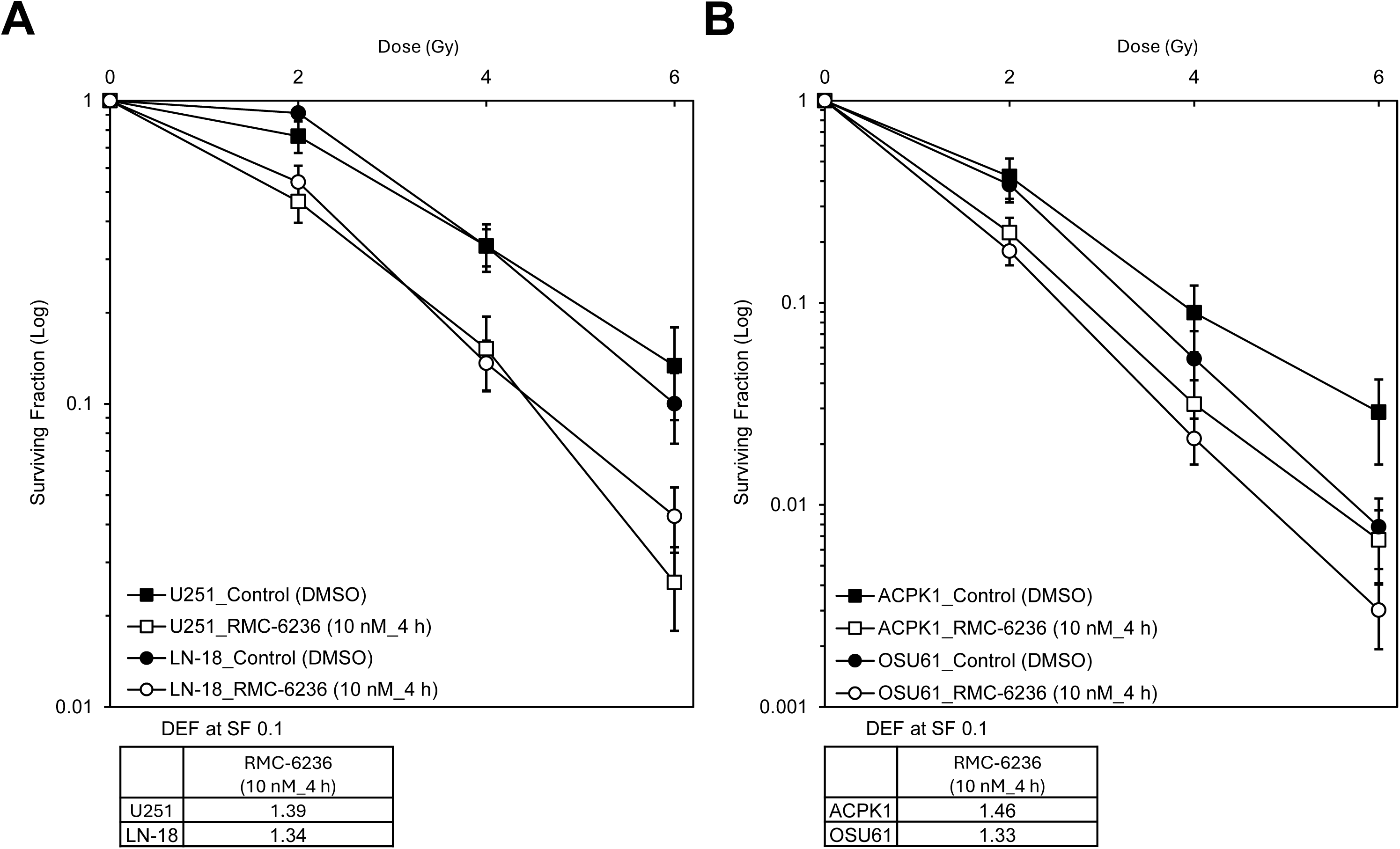
RMC-6236 enhances radiosensitivity in GBM cells. (A-B) Clonogenic survival assays were performed to evaluate the radiosensitizing effect of the KRAS inhibitor RMC-6236 in human glioblastoma (GBM) cell lines. Cells were pretreated with RMC-6236 (10 nmol/L) for 4 h prior to exposure to increasing doses of radiation. (A) Survival curves for U251 and LN-18 cells following treatment with RMC-6236 or DMSO control and irradiation at the indicated doses. (B) Survival curves for ACPK1 and OSU61 cells following treatment with RMC-6236 or DMSO control and irradiation at the indicated doses. The dose enhancement factor (DEF) was calculated at a surviving fraction of 0.1 (SF0.1). The DEF values were 1.39 for U251, 1.34 for LN-18, 1.46 for ACPK1, and 1.33 for OSU61. Data represent mean ± SD from independent experiments. Points, mean; bar, SE. DEF, dose enhancement factor; IR, irradiation; SF, surviving fraction.

### RMC-6236 enhances radiation-induced DNA damage and mitotic catastrophe in GBM cells

To investigate the mechanism underlying RMC-6236–mediated radiosensitization, we examined γH2AX foci formation, a widely used marker of DNA double-strand breaks, and its resolution, which is associated with DNA damage repair (17, 18). We also evaluated mitotic catastrophe, which is a common consequence of unrepaired radiation-induced DNA damage. Four human glioblastoma (GBM) cell lines (U251, LN-18, ACPK1, and OSU61) were pretreated with RMC-6236 (10 nmol/L) for 4 h and subsequently exposed to 4 Gy radiation. γH2AX foci were quantified 1 and 24 h after irradiation. As shown in Fig. 6A and Supplementary Fig. S2-1, RMC-6236 treatment alone did not significantly increase the number of γH2AX foci in any of the four GBM cell lines, indicating that drug treatment alone did not induce detectable DNA damage. Radiation alone resulted in a marked increase in γH2AX foci at 1 h, whereas the number of foci declined by 24 h, which is consistent with the ongoing repair of radiation-induced DNA damage. In contrast, combined treatment with RMC-6236 and radiation led to a significantly greater number of γH2AX foci at 24 h compared to radiation alone in all four GBM cell lines, suggesting that RMC-6236 attenuates the repair of radiation-induced DNA damage. To determine whether the radiosensitizing effect of RMC-6236 was associated with the redistribution of cells into a more radiosensitive phase of the cell cycle, a cell cycle analysis was performed. As shown in Supplementary Fig. S3, treatment with RMC-6236 alone did not produce a consistent change in the proportion of cells in the G0/G1, S, or G2/M phases in any of the GBM cell lines. These findings suggest that RMC-6236–mediated radiosensitization is not attributable to cell cycle redistribution. Because persistent DNA damage can lead to mitotic catastrophe (19), we next assessed the effect of RMC-6236 on radiation-induced mitotic catastrophe using α-tubulin immunostaining. As shown in Fig. 6B and Supplementary Fig. S2-2, treatment with RMC-6236 alone resulted in minimal mitotic catastrophe compared to control cells, whereas radiation alone produced a time-dependent increase. Notably, combined treatment with RMC-6236 and radiation further increased the proportion of cells undergoing mitotic catastrophe at later time points compared with radiation alone in all four GBM cell lines. Together, these results suggest that RMC-6236 enhances the persistence of radiation-induced DNA damage and promotes mitotic catastrophe, providing a mechanistic basis for its radiosensitizing effect in GBM cells.

**Figure 6.**
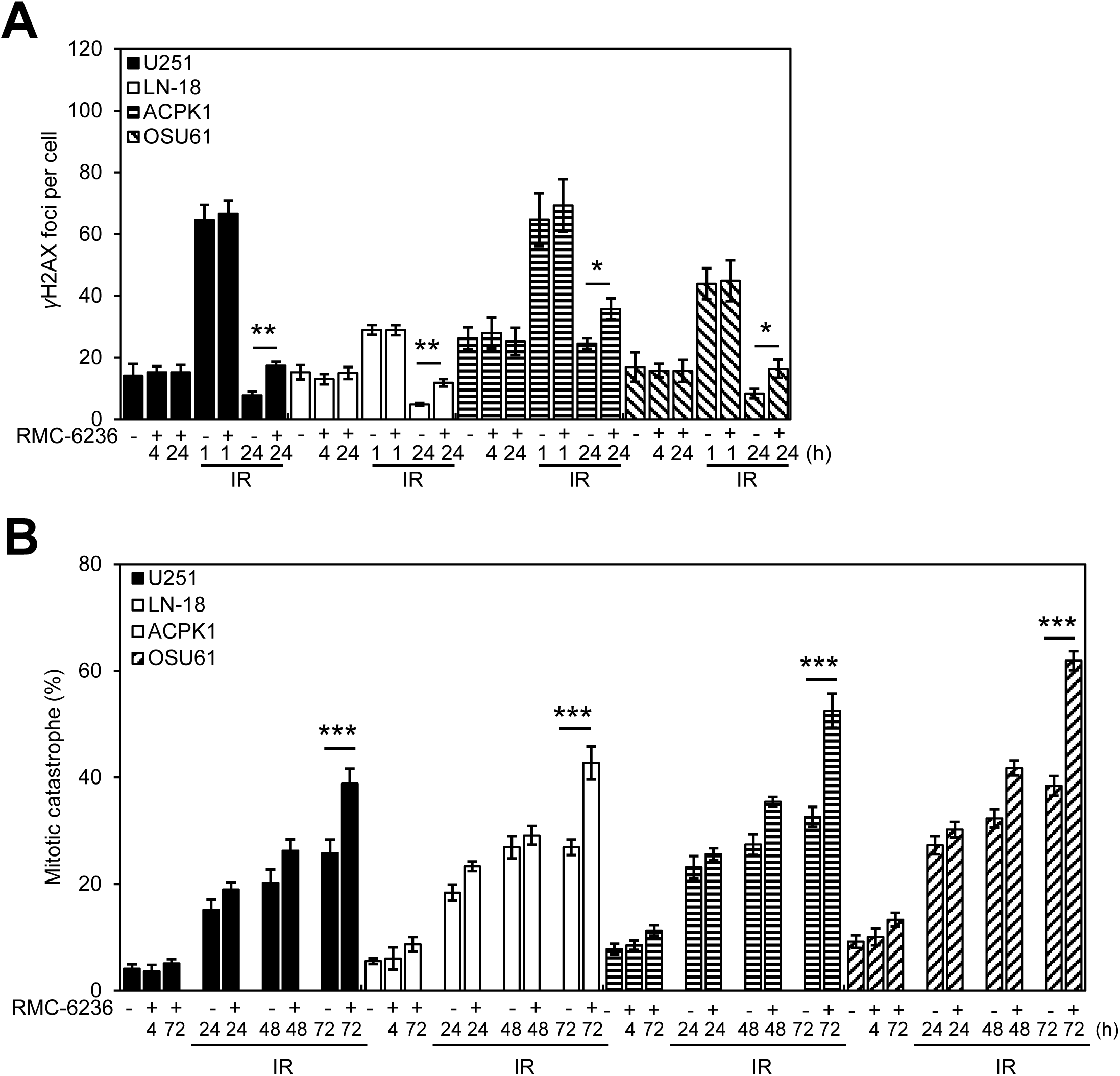
RMC-6236 enhances radiation-induced DNA damage and mitotic catastrophe in GBM cells. Four human glioblastoma (GBM) cell lines (U251, LN-18, ACPK1, and OSU61) were pretreated with RMC-6236 (10 nmol/L) for 4 h prior to exposure to 4 Gy radiation. (A) Quantification of γH2AX foci per cell at 1 h, 4h and 24 h after irradiation. γH2AX foci were scored in at least 50 cells per experiment, and representative histogram images are shown for control cells and cells treated with RMC-6236 alone, radiation alone, or RMC-6236 combined with radiation. (B) Quantification of mitotic catastrophe, expressed as the percentage of cells displaying nuclear fragmentation at the indicated time points following irradiation. Nuclear fragmentation was defined as the presence of two or more distinct nuclear lobes within a single cell, and was evaluated in at least 100 cells per treatment. The data represent the mean ± SEM of three independent experiments. Statistical significance was determined using the Student’s t-test. \**P* < 0.05, \*\**P* < 0.01, \*\*\**P* < 0.001.

### RMC-6236 enhances the radiation response of orthotopic GBM xenografts *in vivo*

To determine whether the radiosensitizing effect of RMC-6236 observed *in vitro* could be translated *in vivo*, we evaluated the therapeutic efficacy of RMC-6236 in combination with radiation using an orthotopic xenograft mouse model established from U251 glioblastoma cells expressing luciferase and GFP (16). Tumor-bearing mice were monitored using bioluminescence imaging (BLI) and randomized into four treatment groups: control (DMSO), RMC-6236 alone, radiation alone, and combination treatment (RMC-6236 + radiation). Kaplan–Meier survival curves were generated to evaluate treatment effects (Fig. 7). Treatment with RMC-6236 alone did not significantly prolong survival compared to that in the control group (P = 0.2014). Similarly, radiation alone failed to produce a significant survival benefit relative to the control (P = 0.3336). In contrast, mice receiving combined treatment with RMC-6236 and radiation exhibited a significant improvement in survival compared to control animals (P = 0.0072). Moreover, combination therapy significantly prolonged survival compared with radiation treatment alone (P = 0.0382). In the Kaplan–Meier curves, statistical significance is indicated by asterisks, where ** denotes the comparison between the control and combination treatment, and * denotes the comparison between radiation and combination treatment. Collectively, these findings demonstrate that, although RMC-6236 or radiation alone had minimal therapeutic effects, the combination of RMC-6236 with radiation significantly improved survival in mice bearing orthotopic U251 glioblastoma tumors, supporting the role of RMC-6236 as a radiosensitizer *in vivo*. The observed therapeutic effect in this orthotopic brain tumor model further suggests that RMC-6236 is capable of exerting antitumor activity within the brain microenvironment, consistent with effective drug delivery across the blood–brain barrier (BBB).

**Figure 7.**
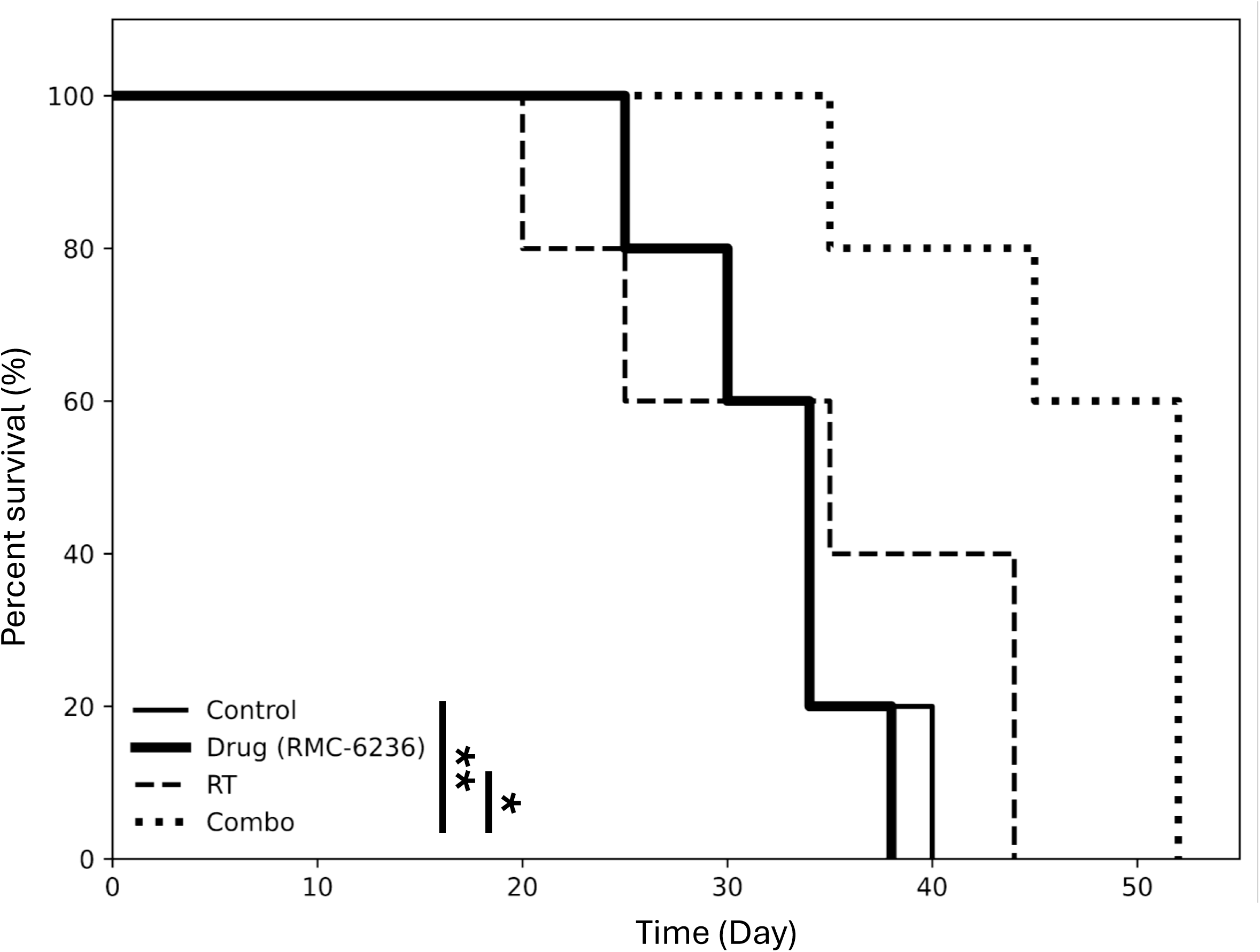
The effect of RMC-6236 on the radiation response of orthotopic GBM xenografts. U251 glioblastoma cells expressing luciferase were implanted orthotopically into mice. On day 6 after tumor implantation, animals were imaged by bioluminescence imaging (BLI) and randomized into four treatment groups: control (vehicle), RMC-6236, radiation, and combination treatment (RMC-6236 + radiation). Mice received 5% DMSO and 95% corn oil or RMC-6236 once daily for three consecutive days, and tumors were locally irradiated (2 Gy) following each drug treatment. This treatment schedule was repeated for three consecutive days. Kaplan–Meier survival curves are shown for each treatment group (control, RMC-6236, radiation, and combination treatment), illustrating the effect of RMC-6236 on the radiation response of orthotopic U251 glioblastoma xenografts. The data represent the mean ± SEM of three independent experiments. Statistical significance was determined using the Student’s t-test. \**P* < 0.05, \*\**P* < 0.01. ** indicates Control vs. Combination treatment, and * indicates radiation vs. combination treatment.

## Discussion

Collectively, our findings support a model in which inhibition of RAS signaling enhances the radiation response in glioblastoma by modulating MAPK signaling, DNA damage persistence, and mitotic catastrophe. Radiation induced rapid activation of KRAS signaling in multiple human GBM cell lines, accompanied by increased phosphorylation of downstream MAPK components, including ERK, p38, and JNK (Fig. 1). Genetic suppression of KRAS using siKRAS inhibited radiation-induced KRAS activation and attenuated downstream MAPK signaling, demonstrating that KRAS is a critical mediator of radiation-induced signaling responses in GBM cells (Fig. 2). Consistent with this observation, KRAS knockdown significantly increased radiosensitivity in GBM cells, supporting the functional role of KRAS signaling in mediating radiation resistance (Fig. 3). Pharmacological inhibition of KRAS signaling using RMC-6236 suppressed radiation-induced KRAS activity and reduced the activation of MAPK signaling pathways, particularly ERK signaling, across multiple GBM cell lines (Fig. 4). In clonogenic survival assays, short-term pretreatment with RMC-6236 significantly enhanced radiation-induced cell death, demonstrating that pharmacological inhibition of RAS signaling can radiosensitize GBM cells (Fig. 5). Mechanistically, combined treatment with RMC-6236 and radiation resulted in persistent DNA damage, as demonstrated by sustained γH2AX foci formation, indicating impaired resolution of radiation-induced DNA double-strand breaks (Fig. 6 and Supplementary data). Consistent with the persistence of DNA damage, RMC-6236 in combination with radiation significantly increased mitotic catastrophe, suggesting that inhibition of RAS signaling compromises DNA damage repair and promotes mitotic failure following irradiation (Fig. 6 and Supplementary data). Importantly, in an orthotopic xenograft model of GBM, combined treatment with RMC-6236 and radiation significantly improved overall survival, supporting the translational relevance of RAS inhibition as a radiosensitizing strategy in vivo (Fig. 7).

Recent advances in RAS-targeted therapeutics have led to the development of mutation-specific KRAS inhibitors that selectively target defined KRAS variants, most notably, KRAS G12C. Agents such as sotorasib and adagrasib have demonstrated significant clinical activity in tumors harboring KRAS G12C mutations, by selectively inhibiting the inactive GDP-bound state of mutant KRAS (8, 9). However, the activity of these agents is inherently restricted to tumors containing the KRAS G12C mutation and does not extend to tumors driven by other KRAS variants or upstream activation of wild-type RAS signaling.

This limitation is particularly relevant to glioblastoma, in which activating KRAS mutations are rare, whereas aberrant RAS–MAPK signaling is more commonly driven by upstream receptor tyrosine kinase activation (12, 20, 21). Analysis of publicly available datasets, including DepMap/CCLE, indicated that the GBM cell lines used in this study, U251 and LN-18, do not harbor hotspot KRAS mutations (22, 23). Consistent with these observations, commonly used GBM cell lines often retain wild-type KRAS, suggesting that RAS pathway activation in glioblastoma more often reflects upstream signaling input than direct KRAS mutations. In this context, therapeutic strategies capable of suppressing both mutant and wild-type RAS signaling may provide broader inhibition of oncogenic signaling networks than mutation-specific KRAS inhibitors (13).

Consistent with this concept, the clinical utility of currently available KRAS inhibitors for glioblastoma appears to be limited. To date, there is little established clinical evidence demonstrating the meaningful antitumor activity of the KRAS G12C inhibitors sotorasib and adagrasib in an unselected GBM population. For sotorasib, the available evidence for GBM appears to be limited to a single case report describing prolonged disease stabilization in a patient with recurrent KRAS G12C-mutated glioblastoma (24). In the case of adagrasib, central nervous system (CNS) activity and cerebrospinal fluid penetration have been reported in patients with KRAS G12C-mutant non-small cell lung cancer (NSCLC) with brain metastases; however, these findings do not provide direct evidence of its clinical efficacy in patients with primary GBM (25). To our knowledge, there are currently no clinical reports demonstrating that either sotorasib or adagrasib has been successfully used as radiosensitizers in patients with GBM. Achieving effective and sustained drug exposure within brain tumors remains a major challenge for molecular targeted therapies for GBM.

In contrast to mutation-restricted KRAS inhibitors, RMC-6236 is an RAS(ON) multi-selective inhibitor designed to suppress active RAS signaling across multiple RAS states, including the wild-type RAS. Although the radiosensitizing activity of RMC-6236 has not yet been clinically validated, accumulating preclinical and translational evidence indicates that this agent exhibits broad antitumor activity in multiple RAS-driven cancers. RMC-6236 demonstrated tumor regression in several solid tumor models, including pancreatic ductal adenocarcinoma, non–small cell lung cancer, and colorectal cancer, and is currently being evaluated in ongoing clinical trials for RAS-mutant cancers (13, 26). Early phase clinical findings have also shown an encouraging safety profile and preliminary antitumor activity in patients with KRAS-mutant solid tumors, including pancreatic and lung cancers. A phase III clinical trial is currently evaluating daraxonrasib (RMC-6236) versus standard therapy for metastatic pancreatic ductal adenocarcinoma (13, 26, 27).

Taken together, our findings suggest that RMC-6236 may overcome the key limitations of mutation-restricted KRAS inhibitors in glioblastoma. By broadly suppressing active RAS signaling, including wild-type RAS, RMC-6236 effectively inhibits radiation-induced MAPK activation and enhances radiosensitivity in GBM models. The improved survival observed in our orthotopic model further suggests that RMC-6236 can exert biological activity within the brain tumor microenvironment. These observations support the concept that pan-RAS pathway inhibition with RMC-6236 represents a promising strategy for enhancing the therapeutic response to radiotherapy in glioblastoma.

## Data availability

All data generated and analyzed in this study are included in the manuscript and its supplementary files.

## Contributions

H.S.Y., A.C., and K.C. designed the study. H.S.Y. and T.K. acquired and analyzed the data. T.K. performed animal experiment. H.S.Y., M.S., K.P, A.C., and K.C. wrote the manuscript, and all authors reviewed the manuscript.

**Supplementary Figure S1-1.**
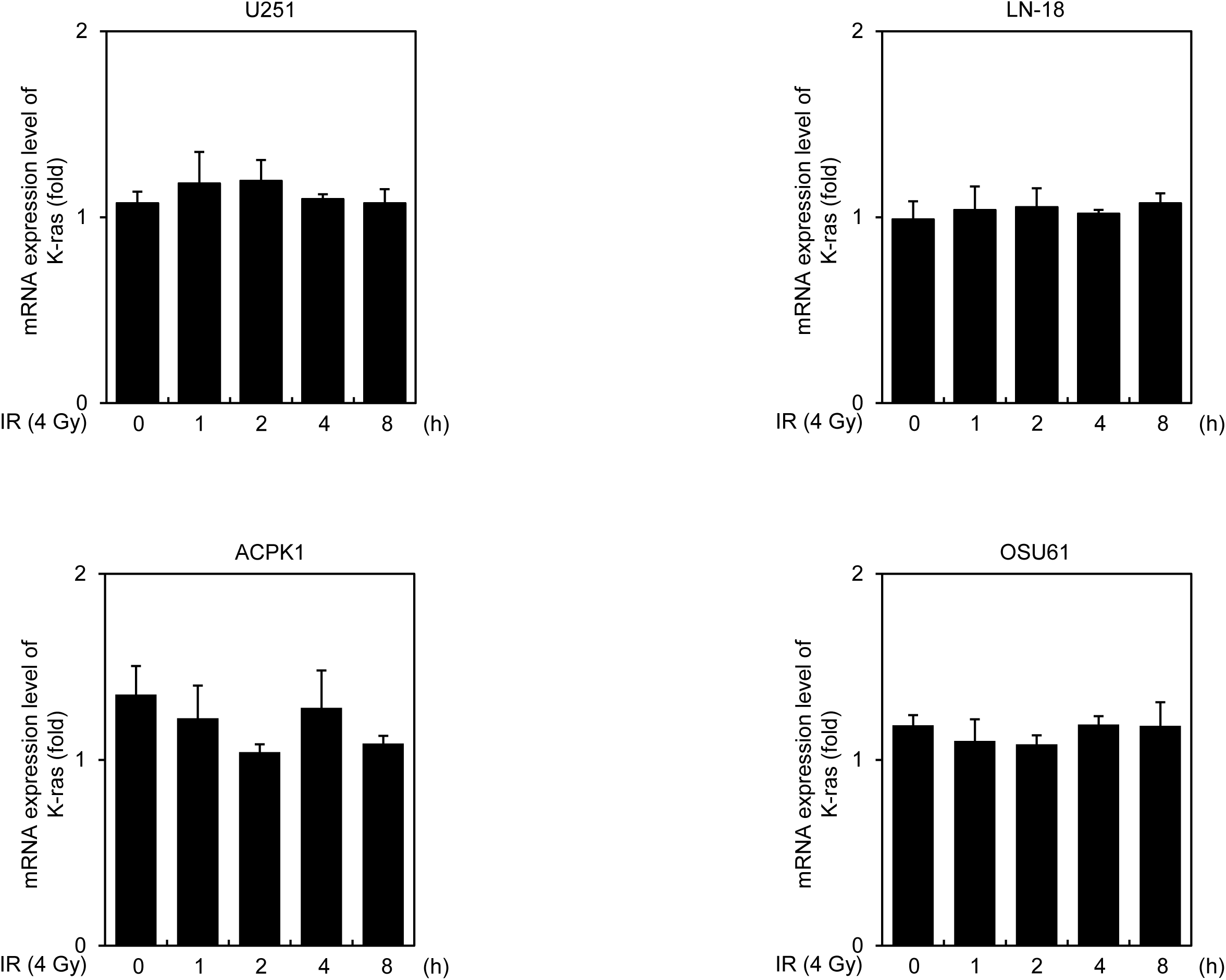
Effect of radiation on KRAS mRNA expression in GBM cells. Quantitative RT-PCR analysis of KRAS mRNA expression in human GBM cell lines (U251, LN-18, ACPK1, and OSU61) following exposure to 4 Gy radiation at the indicated time points.

**Supplementary Figure S1-2.**
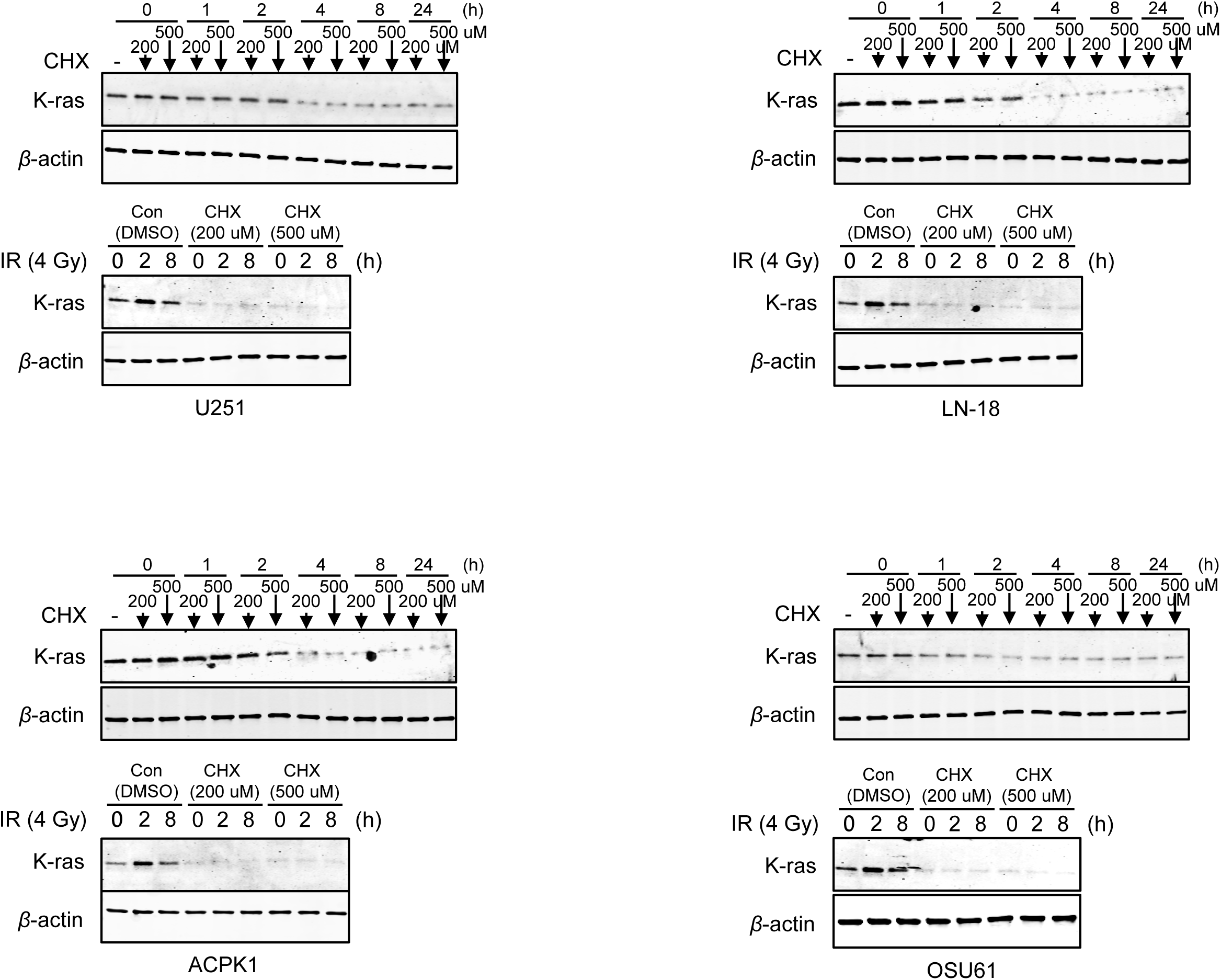
Effect of radiation on KRAS protein stability in GBM cells. Cycloheximide (CHX) chase assays were performed to assess KRAS protein stability. The cells were pretreated with CHX (200 or 500 μmol/L) for 4 h, followed by exposure to 4 Gy radiation. KRAS protein levels were analyzed by immunoblotting at the indicated time points.

**Supplementary Figure S2-1.**
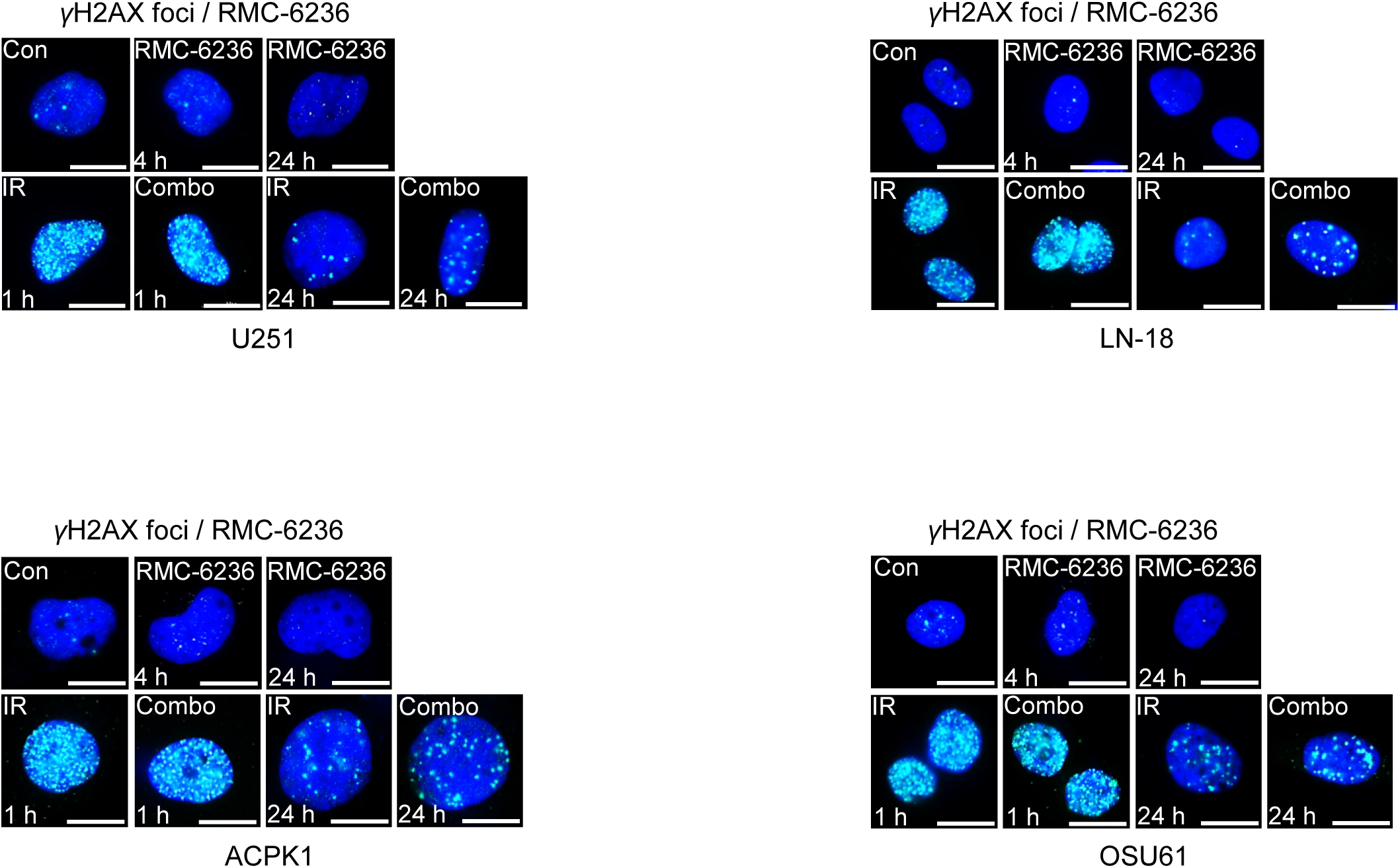
Representative images of radiation-induced γH2AX foci following RMC-6236 treatment in GBM cells. Representative immunofluorescence images of γH2AX foci in U251, LN-18, ACPK1, and OSU61 cells after treatment with RMC-6236 (10 nmol/L) and/or 4 Gy radiation. The cells were analyzed 1 and 24 h after irradiation. Images show control cells and cells treated with RMC-6236 alone, radiation alone, or a combination of RMC-6236 and radiation treatment. Scale bar = 20 μm.

**Supplementary Figure S2-2.**
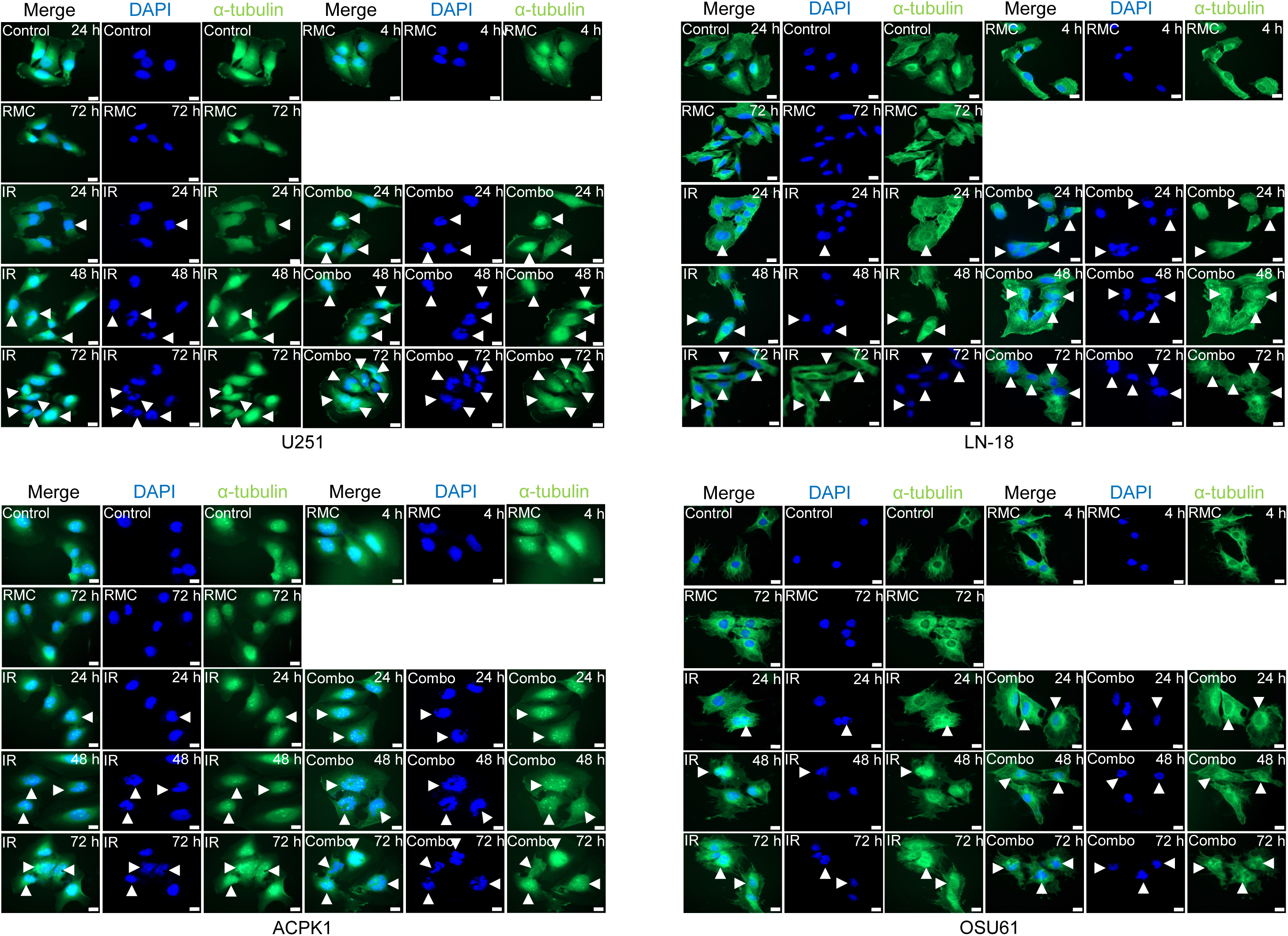
Representative images of mitotic catastrophe following combined RMC-6236 and radiation treatment in GBM cells. Representative immunofluorescence images of mitotic catastrophe in U251, LN-18, ACPK1, and OSU61 cells after treatment with RMC-6236 (10 nmol/L) and/or 4 Gy radiation. The cells were stained for α-tubulin and DAPI and analyzed at the indicated time points after irradiation. Arrowheads indicate cells undergoing mitotic catastrophe characterized by multinucleation or nuclear fragmentation. Scale bar = 20 μm.

**Supplementary Figure S3.**
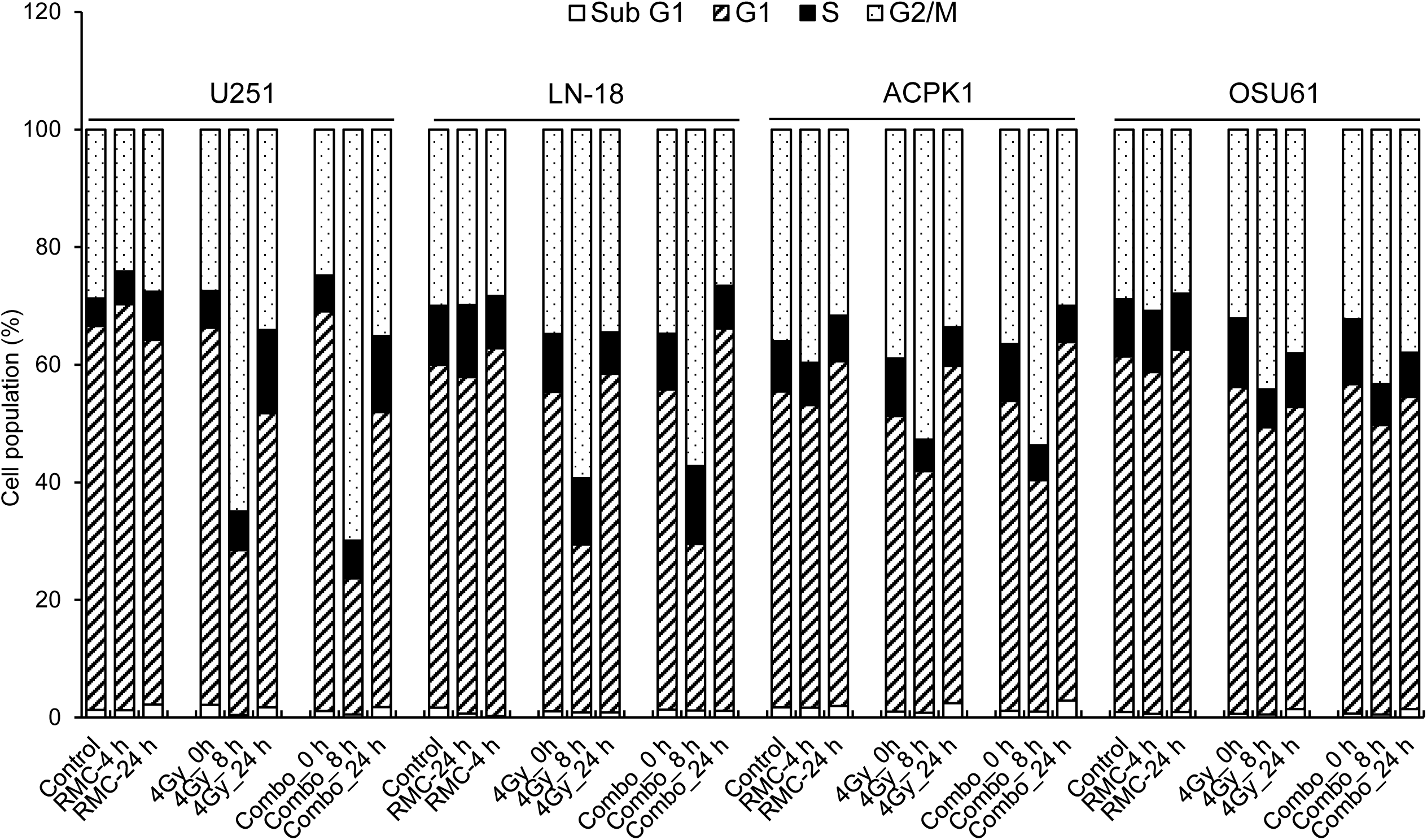
Effect of RMC-6236 on cell cycle distribution in GBM cells. The cell cycle distribution was analyzed in U251, LN-18, ACPK1, and OSU61 cells following treatment with RMC-6236 (10 nmol/L) and/or 4 Gy radiation. The percentage of cells in the G0/G1, S, and G2/M phases was determined by flow cytometry at the indicated time points.

